# Asking the Wrong Questions About American Science Education: Insights from a Longitudinal Study of High School Biotechnology Lab Instruction

**DOI:** 10.1101/2021.11.29.470152

**Authors:** Dave Micklos, Lindsay Barone

## Abstract

The discussion of American science education is often framed by the questions: Why do American precollege students do poorly on international science assessments and what we are doing wrong? Rather we need to ask: Why do so many international students come to US universities for science, what are we doing right in science, and how do we stay ahead in science education? Poor scores on international assessments belie the fact that the U.S. has the best science education system in the world. Our study of 6,200 high school teachers in 1998 and 2018 documented striking success in retooling classrooms for lab-based instruction in biotechnology and provided a pre-COVID-19 snapshot of what is right with American biology education. However, it also highlights the need revitalize our precollege teaching resource with a renewed National Science Foundation commitment to in-service training.

## BACKGROUND

Every several years there is general angst and hand-wringing over American students’ relatively poor performance on the science component of either of two international assessments. U.S. fourth graders ranked 9^th^ and eighth graders ranked 11^th^ in the 2019 Trends in International Math and Science Study (TIMSS)^1^, while 15-year-old students ranked 18^th^ in the 2018 Programme for International Student Assessment (PISA)^2^. These results beg the question: “What is wrong with American science education?”

However, a better question would be, “What do the results actually reflect?” Nine of the top ten scorers on PISA study—Beijing/Shanghai, Singapore, Macao, Estonia, Japan, Finland, Korea, Hong Kong, and Taipei—have centrally-controlled education systems operating in the equivalent of a single large city and/or serving an ethnically homogeneous population. The disparity in the scores of U.S. students reflects both the heterogeneity of the U.S. population and school system—in which 50 state, 13,452 public school districts, and 32,461 private schools exercise local control over curricula^3^. Good scores on standardized tests—international or otherwise—reflect effective test preparation, but do not reflect good preparation for science or support of scientific enterprise.

Heterogeneity may seem a curse on international test performance, but it is a blessing for academic freedom and intellectual curiosity. Despite the U.S.’s lackluster performance on international assessments, the world beats a path to American colleges and universities. In 2019, there were 1,075,406 international students enrolled in U.S. higher education; only about a third as many U.S. students temporarily studied abroad^4^. Foreign students represented 5.5% of total students enrolled in 2019^5^, and 39% of all doctorates awarded in in science and engineering^6^.

Something about our heterogeneous school system works, both in preparing students and in nurturing new scientific ideas. By almost any measure, the U.S. remains the world leader in basic and applied research. Individuals affiliated with U.S. institutions or companies have received 47% of all Nobel Prizes in physics, chemistry, and physiology or medicine^7^ and 51% of all patents awarded by the U.S. Patent and Trademark Office^8^. U.S. scholars were the largest share of top cited authors published in the 2020 H5 citation index of the top five life science journals^9^.

### The Research Edge

Early and frequent student participation in experiments is likely the critical factor in producing good American science. The U.S.’s emphasis on hands-on science means that the better of its heterogeneous schools function as a feeder system for academic and industry research. This science prep system is virtually absent elsewhere in the world—including countries that score well on TIMSS and PISA—as experiments are never an efficient means for rote learning or memorizing facts.

There is strong evidence that participation in research improves Science, Technology, Engineering, and Math (STEM) persistence and academic performance^10^. While the traditional “apprentice” system reaches relatively few hand-picked students, the President’s Council of Advisors on Science and Technology^11^ recommended “replacing standard laboratory courses with discovery-based research courses.” Thus, broadening access to early research experiences has become the holy grail of American collegiate science education, with course-based undergraduate research experiences (CUREs) reaching large numbers of students in the context of for-credit courses. Freshman students who completed a research sequence at the University of Texas at Austin had 23% higher retention in STEM and 17% higher six-year graduation rate than closely matched controls^12^. These effects were consistent across racial groups, and there is mounting evidence for pronounced positive effects of CUREs for underrepresented minorities^13,14,15^. This is significant, because the six-year graduation rate is 23% lower at URM-serving institutions than at mainstream institutions^16,17^. There is every reason to believe that research experiences have similar impacts on high school students. Using the standardized Survey of Undergraduate Research Experiences (SURE-III)^18^, we found that high school students participating in DNA barcoding research scored the same or higher on 12 of 21 learning gains as did college student researchers (Figure 1).

**Figure 1:**
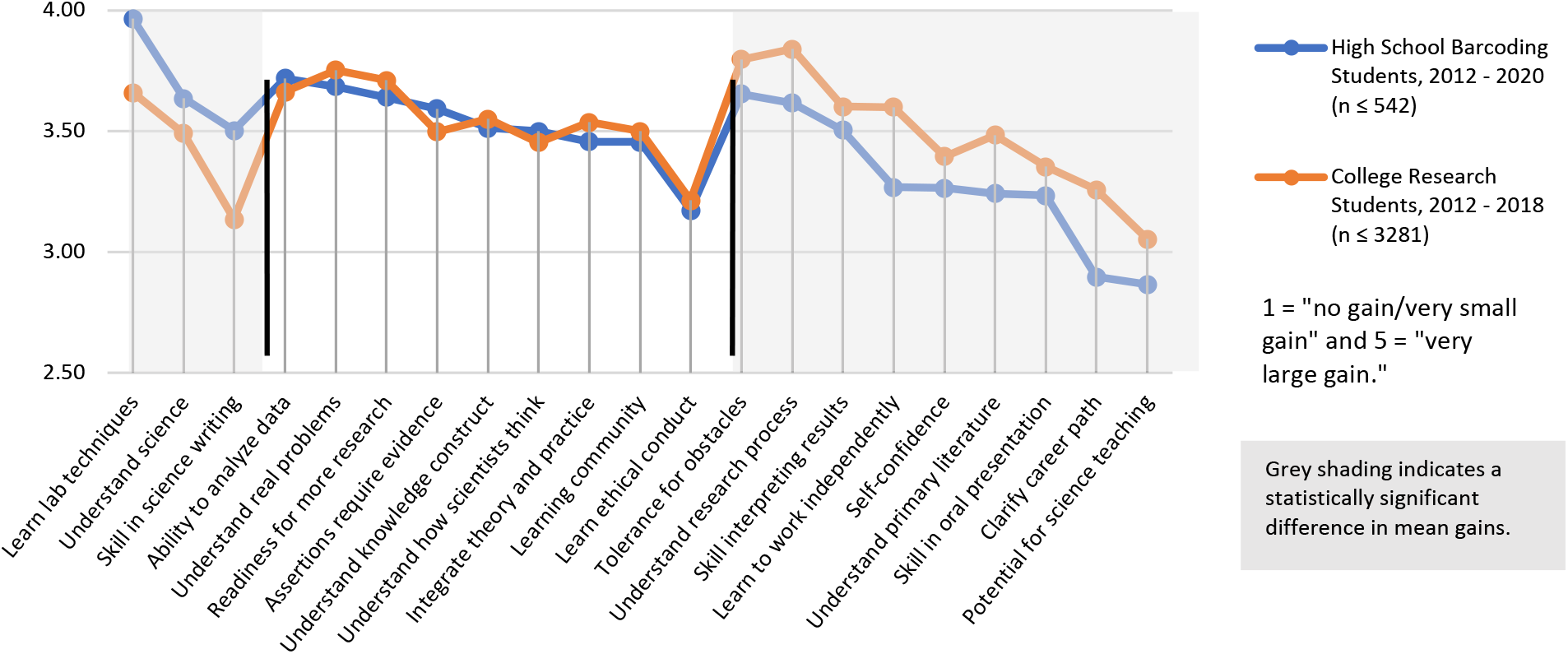
Comparative SURE-III learning gains between high school and college students.

The research experience—identifying a problem, using science tools and methods, persevering to overcome difficulties, collaborating with others, and presenting your work—embodies life skills and a way of looking at the world that is useful beyond science. So, making students ready for research also makes them ready for college, ready for careers, and ready for life. They also learn skills and competencies that prepare them for the jobs of the future.

The best American high schools realize this and support an array of science electives with lab components. Notably, good schools aspire to implement the rigorous college curriculum of Advanced Placement (AP) Biology, which requires students to complete 12 key experiments. Biology is the most popular of all AP sciences (Figure 2), with 260,816 students taking the AP Biology test in 2019^19^.

**Figure 2:**
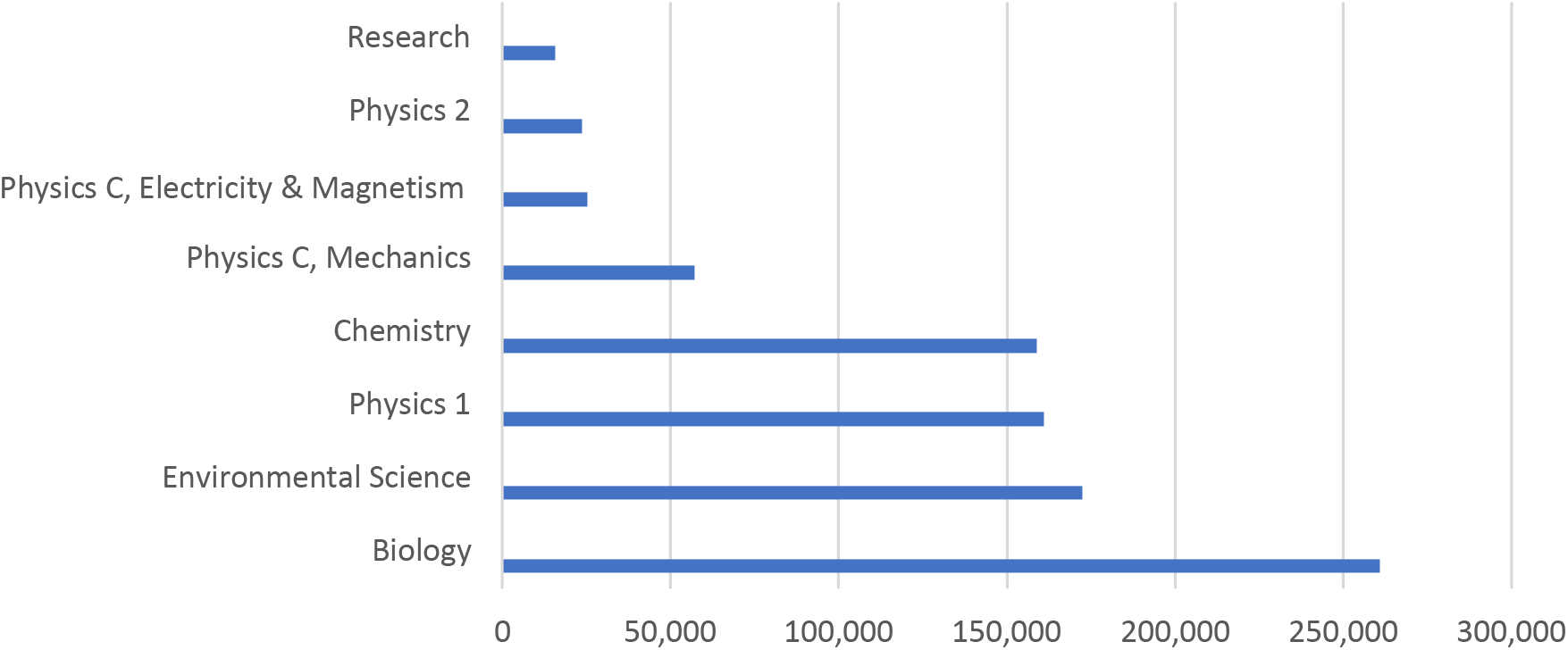
Students taking AP science tests in 2019.

Good schools go well beyond offering AP sciences. For example, virtually all of the 124 school districts on Long Island and many in the New York metropolitan area have high school research programs. These schools accounted for 20% of 2020 finalists in the prestigious Regeneron Science Search^20^. Schools with AP and research programs also tend to infuse inquiry-based labs into their regular science courses. Students from these science-active schools are competitive for admission to the best science universities. They also arrive at college ahead of their peers, ready to participate in early research opportunities— including CUREs that involve an increasing number of freshman students.

### The Biotech Revolution

The DNA Learning Center (DNALC) of Cold Spring Harbor was founded in the mid-1980s, when advances in molecular genetics and biotechnology were changing the face of biology. With funding primarily from the NSF, universities and research institutes stepped in to update high school teachers on new lab techniques. The rapid adoption of hands-on biotech labs in American high schools was catalyzed by the inclusion of two required gene manipulation experiments—DNA restriction analysis and transformation— in the AP Biology curriculum.

The DNALC became a major proponent of hands-on learning in biotechnology. Equipped with our popular lab text *DNA Science* and mobile *Vector* Vans, we could set up cost-effective regional training in molecular genetics at virtually any high school or university. From 1987 to 1996, with NSF and other support, the DNALC instructed 1,931 high school faculty at five-day workshops taught in 39 states. With separate NSF funding, San Francisco State University replicated the *DNA Science* Workshop throughout the state of California—reaching an additional 600 faculty. The *DNA Science* curriculum was also the basis for teacher training workshops administered by the North Carolina Biotechnology Center, University of Wisconsin, and University of Florida that reached 760 participants. All told, over 3,200 high school biology teachers were trained using the *DNA Science* curriculum during this period. Follow-up studies of teachers 17 months after training by the DNALC found that 40% had attempted the two recommended AP Biology labs (Supplemental Table). The rapid uptake of molecular genetics labs in American high schools was also evidenced by rapidly increasing numbers of students who used teaching kits provided by Carolina Biological Supply Company—a major educational science supply house (Figure 3).

**Figure 3:**
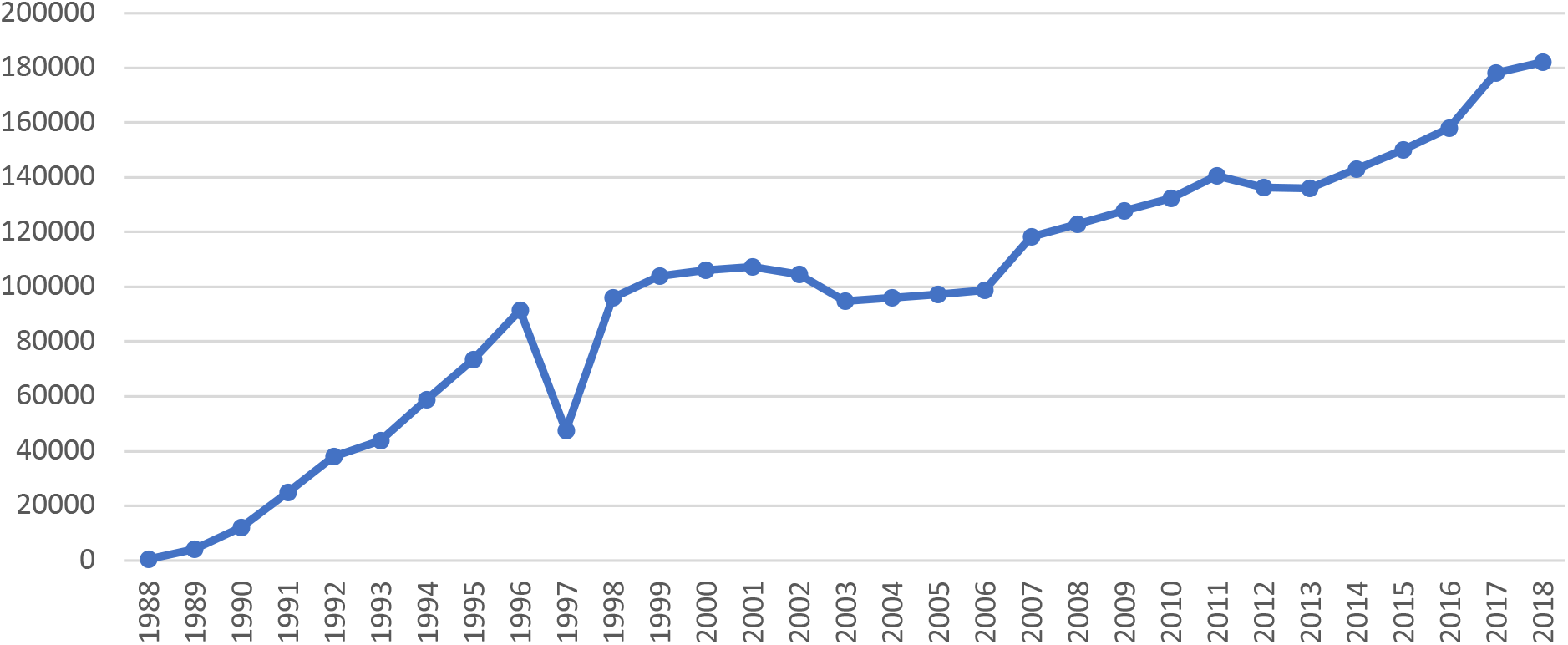
High school students using biotech kits from Carolina Biological Supply Company 1988-2018.

## RESULTS

In 1998, we did a purposive study of 4,100 high school teachers nationwide to gauge the impact of the decade of in-service training done by us and other academic groups. Respondents reported providing 321,826 student exposures to six biotechnology labs in the previous academic year—DNA restriction, transformation, recombination, plasmid isolation, polymerase chain reaction (PCR), and DNA sequencing. Forty-four percent of the teachers reported receiving funding to support biotech instruction, averaging $17,365 each, and 16% of respondents’ schools had started biotech electives. This documented a rapid implementation of new biotech labs and retooling of high school biology classrooms for the gene age.

Responding faculty were highly accomplished and professional educators, with an average of 19 years’ teaching experience in science. Nearly two-thirds had attended at least one professional meeting (65%) and spent at least 16 hours on professional development activities (61%) in the past year. Eighty-one percent were involved in curriculum development, primarily with their own school districts. The respondents were also involved in science-related extracurricular activities—including science field trips (53%), science fairs and competitions (36%), after-school student research (26%), and joint activities with research scientists (24%).

We re-administered the same survey in 2018, to re-take the pulse of biotech lab instruction—this time with 2,100 respondents. We found an equally experienced group of teachers, although slightly younger and with an increased majority of females (69%) (Table 1).

**Table 1:** Teacher demographics, 1998 and 2018.

|  | 1998 | 2018 |
| --- | --- | --- |
| **Average Age** | 52 | 48 |
| **Average Years Teaching** | 19 | 18 |
| **% Fully Certified** | 90% | 93% |
| **% With Graduate Degrees** | 73% | 80% |
| **% Female** | 55% | 69% |
| **% URM in STEM** | 4% | 8% |
| **Average School Size** | 1,165 | 1,285 |
| **Average Class Size** | 24 | 24 |
| **% Teaching AP Biology** | 48% | 37% |

### Key Findings

- **More students participated in biotech labs in 2018. However, fewer faculty were involved in biotech lab teaching, and the pace of integrating new labs slowed.** Scaled for the differing cohort sizes, the number of student exposures to hands-on biotech labs increased 26%, to 406,354. The number of teachers who taught at least one of the labs (in the previous year) decreased from 68% in 1998 to 57% in 2018. Although more 2018 teachers offered newer labs on PCR and DNA sequencing, only a third as many students were exposed to these methods as were to then-current methods of transformation and restriction analysis in 1998 (Table 2).
- **Although fewer 2018 teachers reported funding for biotech programs, per-teacher funding increased by 56%.** Forty-four percent of 1998 teachers received funding to support biotech instruction, averaging $17,365 (adjusted for inflation), while 34% of 2018 teachers received an average of $27,098 in funding.
- **Biotech electives have doubled, but they are not equally distributed and are not well aligned with the school-to-work movement.** Schools offering lab-based biotechnology electives increased from 16% in 1998 to 35% in 2018. However, 64% of schools with electives were located in zip codes above the U.S. median household income. Only 35% of faculty at these schools in 2018 used curriculum materials provided by industry, and even fewer (11%) used materials produced by NSF’s Advanced Technological Education program. Only 22% of these schools had articulation agreements with colleges.
- **Although 2018 faculty were more academically prepared, with 7% more having graduate degrees, they were less involved in professional and extracurricular activities.** Significantly fewer of 2018 teachers belonged to professional societies, including NABT, NSTA, and state science teachers’ associations (Figure 4). While 65% of 1998 teachers had attended one or more professional meetings in the past year, 60% of 2018 teachers had attended none. Significantly fewer 2018 teachers participated with their students in extracurricular activities—including after-school research, science fairs, field trips and joint activities with scientists. Significantly more of the 2018 cohort (41%) said they had done none (Figure 5).
- **Over the last 20 years, teachers continued to believe in-service training is important. However, 2018 teachers perceived that training opportunities have declined.** Teachers in both cohorts rated summer institutes (5 days or more) and workshops at professional meetings as the most important contributors to their own expertise in biotech lab instruction (Figure 6). Thirty-nine percent of 2018 faculty thought there were fewer opportunities for training at workshops and summer institutes than in the past, compared to 27% who think there were more.

**Table 2:** Implementation of six biotechnology labs, 1998 and 2018.

|  | 1998 |  | 2018 |  | n* |
| --- | --- | --- | --- | --- | --- |
|  | % Teaching | Student Exposures | % Teaching | Student Exposures |  |
| **Transformation** | 51% | 80,384 | 45% | 106,453 | 2,101 |
| **Restriction Analysis** | 60% | 118,490 | 43% | 135,240 | 2,437 |
| **DNA Recombination** | 32% | 47,666 | 25% | 65,785 | 1,290 |
| **Plasmid Isolation** | 17% | 24,312 | 15% | 31,044 | 693 |
| **PCR** | 12% | 21,576 | 26% | 28,498 | 559 |
| **DNA Sequencing** | 16% | 29,398 | 20% | 39,334 | 667 |
| **Total Exposures** |  | 321,826 |  | 406,354 |  |

\* Scaled to highest number between the two surveys.

**Figure 4:**
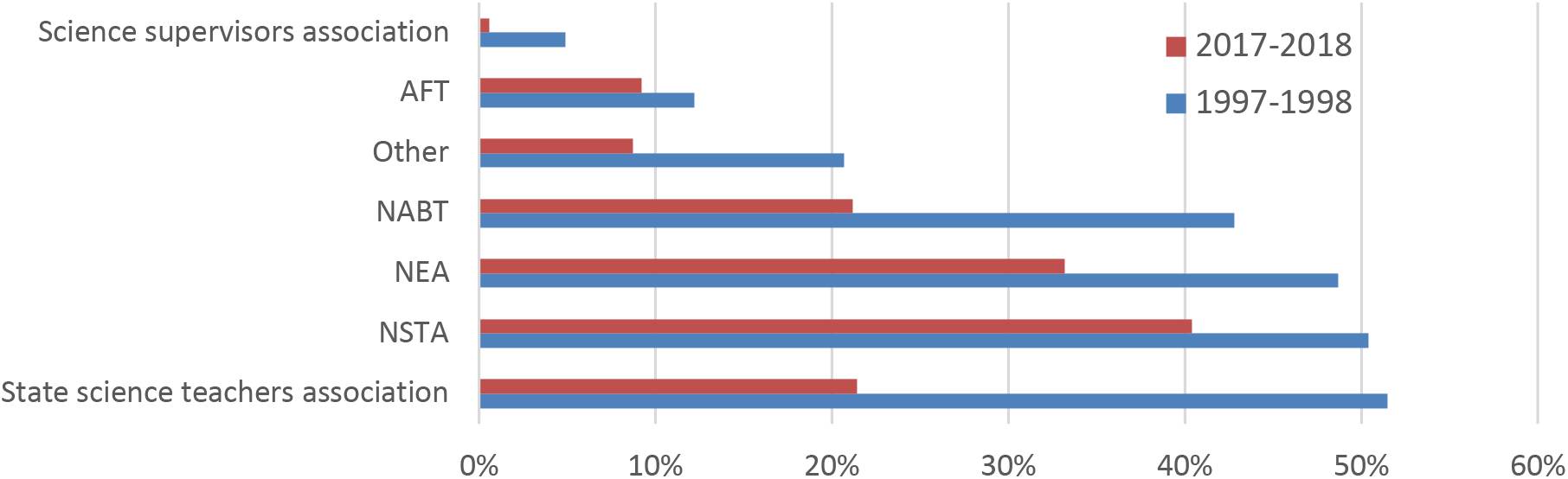
Teacher membership in professional societies, 1998 and 2018.

**Figure 5:**
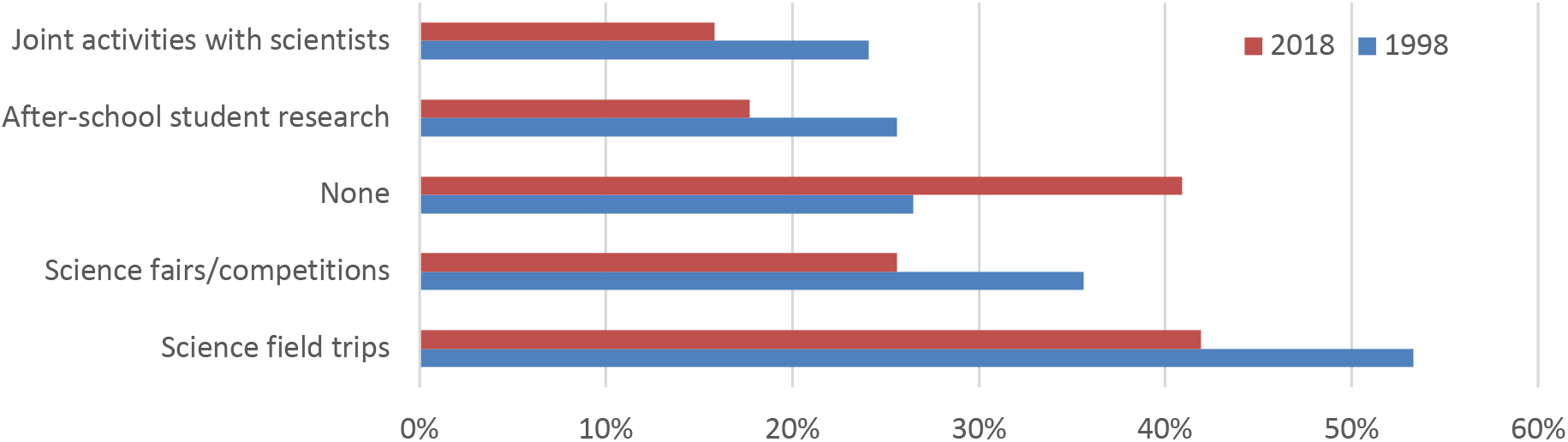
Teacher participation in extracurricular activities, 1998 and 2018

**Figure 6:**
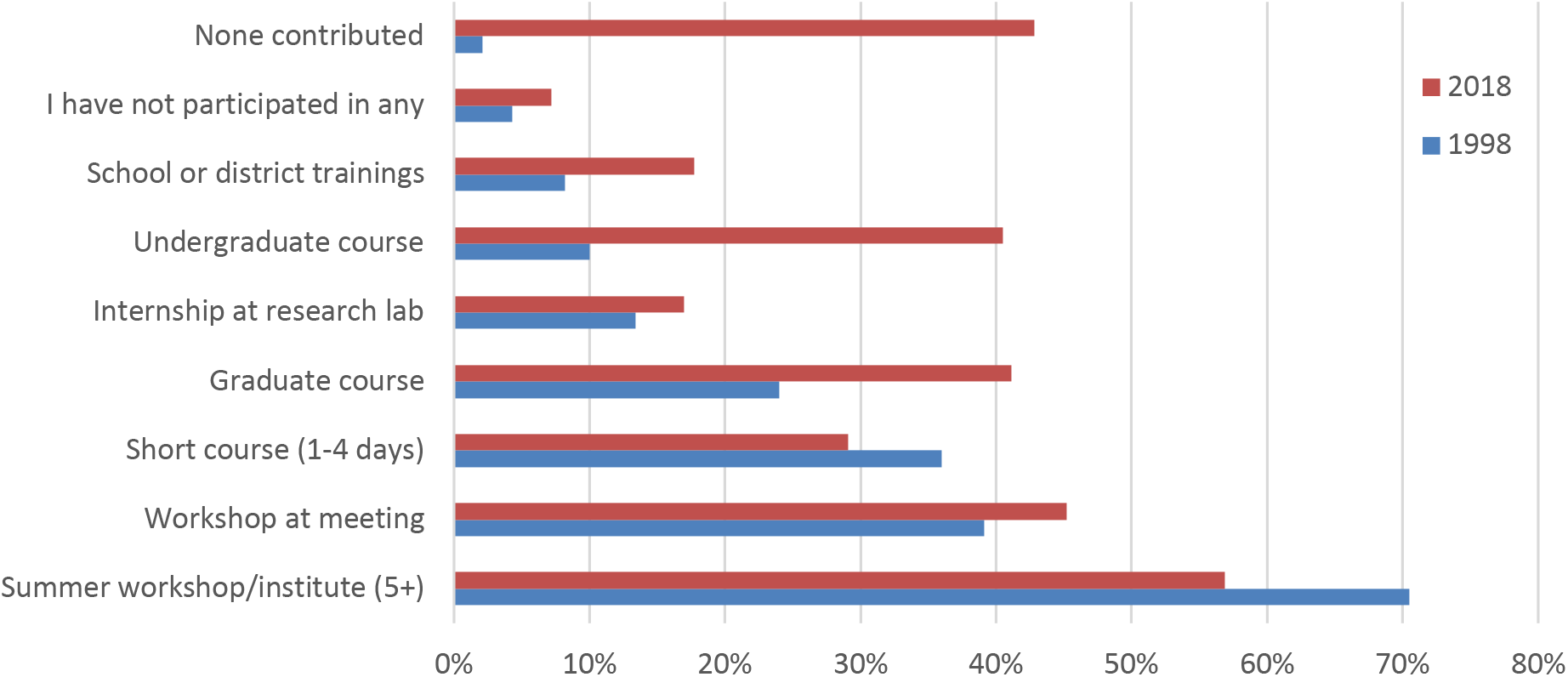
Contributors to teacher innovation in the classroom, 1998 and 2018.

We wondered if this perceived decline was real, so we used keywords to search a database of NSF grants^21^ for potential biology teacher training activities. Searches produced 7,159 unique hits, which we then audited to find 926 projects whose abstracts or titles mentioned any type of training that may have been taken by secondary biology teachers. This included projects in biology, biotechnology, or genetics—as well as general science topics. When we plotted these on a graph, we found that the number of active teacher training grants shrank 75% between its peak in 1994 and 2018 (Figure 7).

**Figure 7:**
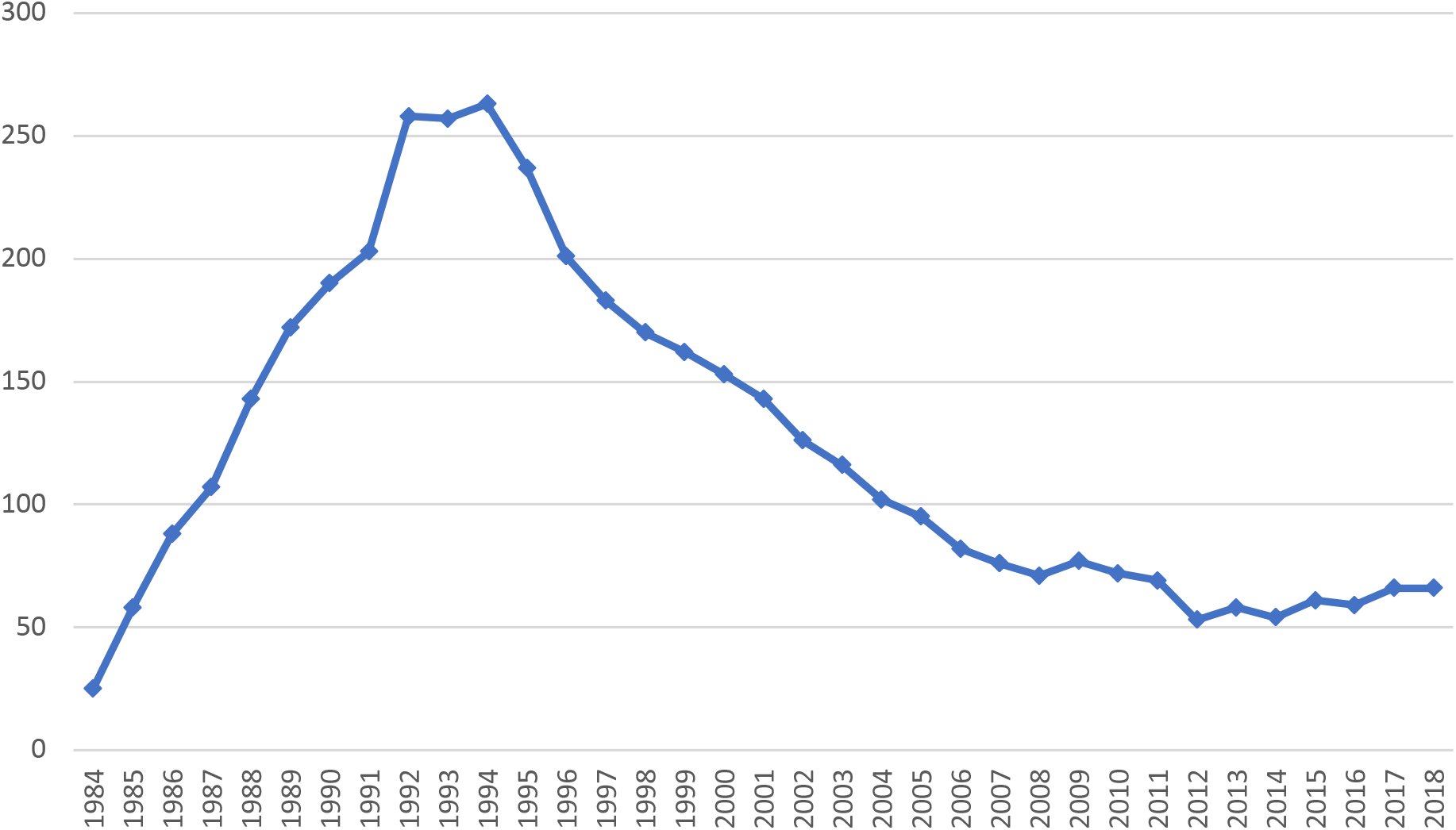
Active NSF science teacher training grants 1984-2018.

## DISCUSSION

The decline in NSF funding for teacher training coincided with an increased emphasis on “systemic initiatives.” A debate about the fundamental value of NSF’s traditional portfolio of teacher-training institutes came to the fore during 1995 meetings of a National Academy of Sciences committee tasked to examine in-service training for high school biology teachers^22^. A vocal minority of the committee criticized traditional NSF summer institutes as “pulling out” the best teachers and leaving the rest behind. The NSF programs of systemic reform, which began in 1990, seemed a logical antidote. These shifted the responsibility for in-service training from content experts at universities and research institutes to state and local school authorities. The content experts had hewed to the NSF priority for innovation, which favored new advancements and experimentation. Local school systems focused on pedagogy and content aimed at “all” teachers. This local training was not necessarily well received by teachers accustomed to the authority and rigor of their previous experiences at NSF summer institutes. A review of 25 state systemic initiatives by SRI International found that professional development activities reached only about 20% of math and science teachers, even though they consumed about one-third of funding. It concluded that “there are few, if any, places that are models of ‘systemic’ professional development, i.e. involving all teachers in high-quality learning on a routine basis…”^23^

The final blow to NSF’s commitment to in-service teacher training came in 2012 with the cancellation of the Teacher Enhancement Program, which had funded the majority of summer institutes that flourished in the 1980s and 1990s. Since then, the Division of Research and Learning has devoted the lion’s share of pre-college education funding to research on the education process. Under this regime, in-service teacher training can only be funded as a component of an educational research program—it is valueless on its own.

After mobilizing scientific research during WWII, Vannevar Bush imagined an “endless frontier”^24^ for American science and provided a template for the National Science Foundation Authorization Act of 1950^25^. NSF sponsorship of summer institutes for secondary science and math teachers began in 1954. By 1965, approximately 146,000 teachers had received training at more than 2,500 institutes, typically of 6-8 weeks’ duration^26,27^. Support for teacher institutes waned in the 1970s and reached a nadir during in 1982, during the first Reagan administration. The advent of the Teacher Enhancement Program in 1984, marked a return to NSF’s tradition of supporting summer science, albeit with shorter and more costeffective institutes of 1-3 weeks. This program nurtured the biology teachers who responded to the challenge of bringing advances in biotechnology into American classrooms.

That generation of biology pioneers has retired, and the biotech revolution they created in American biology classrooms has lost momentum. Little has been done to rebuild their culture of lifelong learning. Our study suggests that teachers’ professionalism and extracurricular involvement with students contracted along with NSF’s commitment to in-service training. Furthermore, lab instruction shrunk to near zero nationwide during the COVID-19 pandemic. As we emerge from a year of isolation, biology teachers will be anxious for opportunities to re-connect with science and their peers. We need to ask, “How we can we help science teachers return to research-driven instruction and to keep up with quickening advances in biological sciences?” The answer is clear.

It is the moment for NSF to take the lead in re-establishing an “esprit de corps” and culture of lifelong learning among American science teachers. NSF must renew its commitment to in-service training as a critical contributor to teachers’ professional development and classroom innovation. Now is the time for every person concerned with science and science education in this country to send two clear messages to policy makers:

1. Reinstate NSF’s mandate for teacher training, and implement an extensive program of summer institutes and academic-year workshops at professional meetings.
2. Provide state and local support for teacher training and attendance at professional meetings.

## ACKNOWLEDGEMENTS

Our thanks to:

- John Kruper, who helped plan the initial teacher study, based on insights from his doctoral research on early cohorts of teacher trainees.
- DNALC staff members who designed and administered the national teacher surveys at two time points—Susan Lauter, Judy Cumella-Korabik, and Valerie Meszaros—and who trained teachers under NSF grants 1986-2018—Mark Bloom, Uwe Hilgert, and Bruce Nash.
- Linnea Fletcher and Celeste Carter, who provided valuable feedback throughout the study.

This research was supported by grants from the National Science Foundation’s Advanced Technological Education (ATE) Program: DUE-1744708, DUE-1839544, and DUE-1901984.

---

**SUPPLEMENTAL TABLE:** Impacts of DNALC Teacher Training 1987-1995

|  | 1987-88 | 1989-90 | 1991-92 | 1993-95 | TOTALS |
| --- | --- | --- | --- | --- | --- |
| Participants* | 443 | 242 | 315 | 201 | 1201 |
| Participants who returned follow-up ** | 390 (88%) | 211 (87%) | 266 (84%) | 176 (88%) | 1043 (87%) |
| **Demographics\*** |  |  |  |  |  |
| Male-Female | 45%-55% | 40%-60% | 35%-65% | 46%-54% | 42%-58% |
| Caucasian | 93% | 90% | 74% | 78% | 85% |
| Public day school | 82% | 83% | 86% | 90% | 85% |
| Median school enrollment | 1,000-1,500 | 1,000-1,500 | 1,000-1,500 | 1,000-1,500 | 1,000-1,500 |
| AP, honors, or elective teaching assignment | 80% | 80% | 78% | 80% | 79% |
| Years teaching biology or science | 16-20 | 16-20 | 11-15 | 11-15 | 16-20 |
| Masters degree or higher | 63% | 74% | 54% | 67% | 64% |
| **Professional Development\*** |  |  |  |  |  |
| NSTA member | 48% | 50% | 46% | 44% | 47% |
| State science teachers organization | 46% | 56% | 50% | 47% | 49% |
| NABT member | 40% | 44% | 31% | 33% | 37% |
| Hours spent on professional development/year | 16-35 | 16-35 | 16-35 | 16-35 | 16-35 |
| Professional conferences attended/year |  | 2 | 3 | 2 | 2 |
| Involved in curriculum/syllabus development | 73% | 70% | 68% | 85% | 73% |
| **Extracurricular Activities\*** |  |  |  |  |  |
| Science field trips | 58% | 62% | 57% | 67% | 60% |
| Science fairs/competitions | 45% | 56% | 56% | 51% | 51% |
| After-school student research | 34% | 38% | 41% | 36% | 37% |
| Joint activities with local scientists | 25% | 28% | 31% | 37% | 29% |
| **Teaching Behavior \*\*** |  |  |  |  |  |
| Gave personal account of recombinant DNA | 94% | 96% | 91% | 91% | 93% |
| Used new examples for topics already on syllabus | 73% | 79% | 72% | 78% | 75% |
| Presented new topics not currently on syllabus | 67% | 73% | 63% | 69% | 68% |
| Labs attempted Basic bacteriology | 66% | 74% | 72% | 73% | 70% |
| Bacterial transformation | 35% | 51% | 46% | 44% | 43% |
| DNA restriction analysis | 25% | 42% | 43% | 54% | 38% |
| DNA recombination | 18% | 27% | 30% | 30% | 25% |
| Plasmid isolation | 13% | 19% | 19% | 18% | 17% |
| **Networking Behavior\*\*** |  |  |  |  |  |
| Made presentation to local educators | 36% | 37% | 32% | 35% | 35% |
| Demonstrated lab to educators | 18% | 18% | 17% | 27% | 19% |
| Led teachers training workshop | 9% | 10% | 8% | 14% | 10% |
| Made presentation to professional organization | 7% | 10% | 8% | 11% | 9% |
| **Funding for Molecular Genetics Instruction\*\*** |  |  |  |  |  |
| Cash funding | \$381,000 | \$291,000 | \$364,000 | \$598,050 | \$1,634,050 |
| Equipment/supplies donation value | \$126,000 | \$203,000 | \$177,000 | \$181,055 | \$687,055 |
| Total funding | \$507,000 | \$494,000 | \$541,000 | \$779,105 | \$2,321,105 |
| Per-participant funding | \$1,144 | \$2,041 | \$1,717 | \$3,876 | \$1,933 |

\*Measured at time of DNA Science training.
\*\*Measured 17 months after training.

